# Seizures start as silent microseizures by neuronal ensembles

**DOI:** 10.1101/358903

**Authors:** Michael Wenzel, Jordan P. Hamm, Darcy S. Peterka, Rafael MD Yuste

## Abstract

Understanding seizure formation and spread remains a critical goal of epilepsy research. While many studies have documented seizure spread, it remains mysterious how they start. We used fast in-vivo two-photon calcium imaging to reconstruct, at cellular resolution, the dynamics of focal cortical seizures as they emerge in epileptic foci (intrafocal), and subsequently propagate (extrafocal). We find that seizures start as intrafocal coactivation of small numbers of neurons (ensembles), which are electrographically silent. These silent “microseizures” expand saltatorily until they break into neighboring cortex, where they progress smoothly and first become detectable by LFP. Surprisingly, we find spatially heterogeneous calcium dynamics of local PV interneuron sub-populations, which rules out a simple role of inhibitory neurons during seizures. We propose a two-step model for the circuit mechanisms of focal seizures, where neuronal ensembles first generate a silent microseizure, followed by widespread neural activation in a travelling wave, which is then detected electrophysiologically.

## Introduction

Understanding the properties of seizure-producing cortical circuits (“ictal networks”) may enable more efficient seizure control. Epileptic discharges are thought to emerge at locally confined epileptic foci, from where they can secondarily spread across the brain. Although there have been several studies detailing how epileptic seizures invade and spread through neighboring cortical territories (Badea et al., 2001; Cammarota et al., 2013; Feldt Muldoon et al., 2013; Lillis et al., 2015; Muldoon et al., 2015; Neubauer et al., 2014; Tashiro et al., 2002; Trevelyan et al., 2007; Wenzel et al., 2017), it remains unclear how seizures actually start *in vivo*. Further, because cortical circuits are complex in terms of cell types and connectivity, it is likely that the contribution of individual cell types to seizure progression could differ depending on their location. Yet to date, little is known about the local neural sub-population dynamics within ictal networks (Muldoon et al., 2015; Neumann et al., 2017). However, an understanding of how exactly seizures progress from micro- to macroscale in the intact brain may hold critical new clues on how to stop their expansion.

Due mostly to technical limitations, our knowledge about the precise dynamics of local brain networks in and outside such foci is still scarce. Recent technologies have enabled more fine-scaled studies of these dynamics leading for example to the identification of micro-epileptic discharges (“microseizures”) (Goldensohn, 1975; Schevon et al., 2008; Stead et al., 2010; Worrell et al., 2008), which are clinically silent and too small and confined to be detectable by conventional electrodes (Stead et al., 2010). Thus, a better understanding of seizure emergence and spread at a neural circuit level requires techniques with high spatiotemporal resolution. While multi-electrode arrays have pushed our understanding of fine-scale epileptic network dynamics, the most common arrays (96 electrodes, 4×4mm) do not allow measurements of complete neural circuits. In fact, such arrays usually record from several dozen (at best up to ∼180) neurons from patches of cortex or hippocampus (Neumann et al., 2017; Truccolo et al., 2011), which, however, contain hundreds of thousands of neurons. Due to this, and the usually spatially poor identification of seizure initiation sites prior to electrode implantation in patients or animals, it has remained unproven if micro-seizures are mandatory pre-cursors of pending macro-seizures, or how they transition from focal discharges to generalized ictal events. As an alternative to electrophysiology, calcium imaging can monitor action potential activity with single cell resolution (Smetters et al., 1999; Yuste and Katz, 1991), so it seems as an ideal method to map microseizures and ictal spread, cell by cell (Badea et al., 2001). However, calcium imaging studies of epileptiform activity have traditionally suffered from a temporal resolution too low to investigate sizable population dynamics during seizures (Badea et al., 2001; Cammarota et al., 2013; Feldt Muldoon et al., 2013; Lillis et al., 2015; Muldoon et al., 2015; Neubauer et al., 2014; Tashiro et al., 2002; Trevelyan et al., 2007). To improve the temporal resolution of calcium imaging, we recently introduced a fast (30Hz) resonant scanning resonant scanning method to study cortical seizure propagation in vivo (Wenzel et al., 2017). In that initial study, general local network recruitment patterns to spreading ictal activity were investigated at a distance to the seizure initiation site. We found that in the propagation area, seizure spread occurs smoothly, through stereotypical circuits, and that this spread is elastic in time and regulated by the activity of GABAergic interneurons.

Here we turn our attention to the initiation site of cortical seizures (‘epileptic focus’ or ‘intrafocal region’), and compare the findings to recruitment patterns within the propagation area (‘surround cortex’ or ‘extrafocal region’). To compare seizure progression in both these compartments, we use fast (30Hz) two-photon calcium imaging of GCaMP-labeled neuronal populations of mouse somatosensory cortex and LFP recordings in the 4-Aminopyridine (4-AP) mouse model of local onset seizures *in vivo*. The approach combines high temporal and single cell resolution to unveil the local population activity underlying seizures with unequivocal anatomical precision. With a precisely defined seizure onset zone by local cortical 4-AP injection, we image cortical circuits within the seizure initiation site as well as the propagation area that is invaded only during ictal spread. Further, we perform sub-population calcium imaging of td-Tomato labeled GABAergic interneurons (parvalbumin-positive or PV neurons) within those territories during seizure emergence and spread, and find spatially heterogeneous PV recruitment, in contrast to simple models of how GABAergic interneurons are recruited during epilepsy. Our data represent a complete reconstruction of seizures progression from their microepileptic origins, have potential implications for the early detection of pending seizures, and provide novel insights into the recruitment dynamics of neuronal sub-populations in neocortical focal onset seizures.

## Results

We used an *in vivo* mouse model of acute cortical seizures applying local injection of 4-AP (15mM, 500nl, layer 4 or 5, total amount delivered = 7,5nmol) in lightly anesthetized mice (n=13 animals). In six mice we characterized the intra- or extrafocal recruitment to epileptic activity (898 neurons), while seven animals were used for imaging of neuronal sub-populations (1059 neurons, of which 79 were PVs). We carried out experiments under light anesthesia to reduce the animal burden of a series of prolonged tonic-clonic seizures. Despite differences such as seizure threshold or propagation speed, general neural recruitment patterns during cortical seizure progression share fundamental characteristics across anesthesia and wakefulness (Uva et al., 2013; Wenzel et al., 2017). Indeed, seizures in humans do not only happen during wakefulness but are encountered in seizure-susceptible individuals even during deeper anesthesia on the intensive care unit or in the operation room (Howe et al., 2016; Ulkatan et al., 2017). The 4-AP model was chosen for several reasons (see also Methods). First, local injection of 4-AP establishes a territorially well-defined acute epileptic focus surrounded by otherwise unperturbed cortex that is invaded during seizure spread. This acute situation actually resembles real world medical conditions such as intracerebral bleeding, local brain trauma or ischemic stroke, in which acute seizures often occur shortly after the injury (Beleza, 2012). The model also shares similarities with chronic conditions such as epilepsy with focal onset seizures, where spatially confined epileptic discharges secondarily spread into otherwise functionally intact brain tissue (Milton et al., 2007). Moreover, unlike disinhibitory chemoconvulsants like picrotoxin or bicuculline, 4-AP leaves inhibitory circuits intact, and elicits electrographic and behavioral phenomena that are similar to seizures in chronic epilepsy (Avoli et al., 2002; Szente and Pongracz, 1979). Instead of addressing a specific disease pathway but rather considering epileptic seizures as a phenomenon shared by many neurological disorders and even the healthy brain, our model focused on understanding the phenomena of epileptic states (Jirsa et al., 2014).

### Two-photon calcium imaging of ictal foci *in vivo*

Two experimental setups were employed: either a craniotomy for imaging within the propagation area or a craniotomy combined with adjacently thinned skull for intra-focal imaging (Fig. 1 A). In both setups, imaging was carried out in layers 2/3 (L2/3, ∼150 µm beneath the cortical surface) of mouse primary somatosensory cortex. The experimental workflow is depicted in Figure 1 B. For LFP measurements, which served as a gross indicator and additional confirmation of epileptic activity within the investigated cortical territory, a sharp glass microelectrode was carefully lowered into the cortex close by the imaged area. For induction of ictal events, a second glass micropipette containing 4-AP (total amount delivered = 7,5nmol) was slowly advanced to a cortical depth of ∼480 µm. The tip of the 4-AP pipette was either positioned at a distance of ∼1,5-2 mm caudally to the field of view (propagation area) or directly underneath the imaged area (initiation site). During baseline conditions, local populations in L2/3 in the epileptic focus displayed ongoing sparse and distributed calcium activity (Fig. 1 C left, 1 D left; (Miller et al., 2014)), contrasted by sustained firing of larger numbers of neurons in the field of view during seizure occurrence post 4-AP injection (Fig. 1 C right, 1 D right). Within the surrounding cortex, ictal invasion happened in a wave of burst neuronal firing continuously advancing across the field of view (suppl. movie 1). The temporal imaging resolution of 30Hz was sufficient to capture individual cell recruitment to ictal activity (Fig. 1 E). To better understand this recruitment, we focused on the immediate pre-ictal and early intra-ictal cell recruitment (time window 45 seconds prior to and 10 seconds into individual seizures), since this window represents a critical time for potential therapeutic interventions. In order to systematically investigate general temporal characteristics of seizure progression, we analyzed the activity of all neurons in the field of view that showed visible calcium transients during epileptic activity and whose somata could be followed across the entire experiment. Individual cell recruitment time-points to ictal activity were derived by calculating the first discrete derivative (slope) of the fluorescent traces assuming that the steepest rise in fluorescence correlates best with maximal recruitment to seizure activity (Badea et al., 2001; Trevelyan et al., 2006; Wenzel et al., 2017).

**Figure 1:**
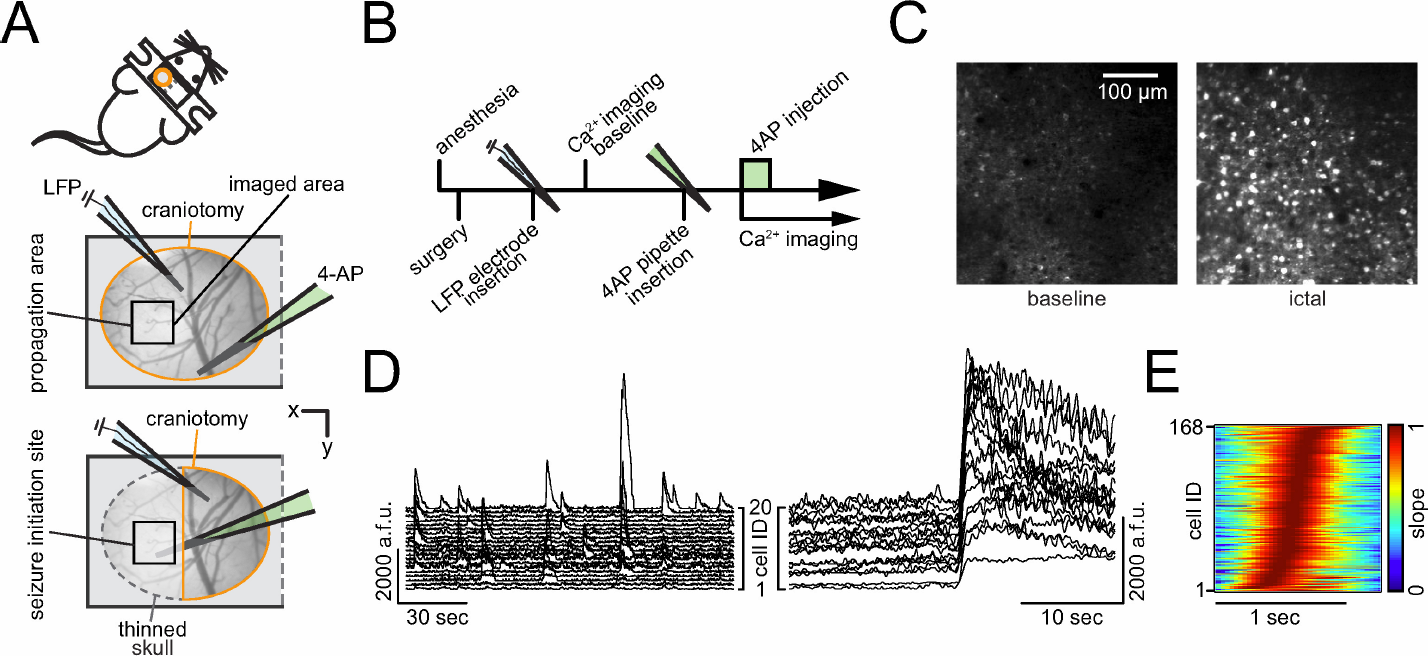
**Imaging seizures in initiation and propagation area** (**A**) Experimental setup. Two surgical approaches over left somatosensory cortex; craniotomy encircled in orange, thinned skull in dotted gray, black squares indicate imaging field of view (FOV); each experiment (exp.) involved the insertion of two glass micropipettes, one (blue) containing a silver chloride silver for LFP (local field potential) recording, the other (green) containing 4-AP (4-Aminopyridine, 15mM, injection vol. 500 nl, total amount delivered = 7,5nmol). (**B**) Experimental workflow. (**C**) Propagation area, representative 3 second average fluorescence images of neural activity (GCaMP6s) during baseline and ictal event. (**D**) Calcium transients of 20 representative cells during baseline conditions (left) and during seizure onset period (right). a.f.u. = arbitrary fluorescent units (**E**) Propagation area, representative example of the arriving ictal wavefront. Normalized first derivative of each registered neuron’s fluorescent trace during electrographic seizure onset. Cell recruitment to ictal activity ordered in time by maximum slope. Note the s-curved shape of cell recruitment highlighting sufficient temporal imaging resolution for individual cellular recruitment.

### Seizure initiation and propagation areas display differential spatiotemporal activity

In the propagation area, electrographic seizures and calcium transients corresponded well and neurons in the propagation area were recruited in a continuous fashion (Fig. 2 A, Fig. 2 B top) (Wenzel et al., 2017). But, prior to ictal invasion during electrographic seizure onset, little to no calcium activity was visible in surround cortex. To analyze this, we superimposed the calcium activity for all experiments in the propagation area around the frame where the proportion of active neurons reached 50%. The superimposed graphs lined up consistently, describing a continuous s-shaped curve (Fig. 2 B bottom, n = 3 experiments; GCaMP6s; total # of seizures = 31; average # of seizures = 10,3 ± 1,5275; total # of analyzed cells = 576; average # of analyzed cells = 192 ± 58,9237), as previously described (Neumann et al., 2017; Wenzel et al.,2017).

**Figure 2:**
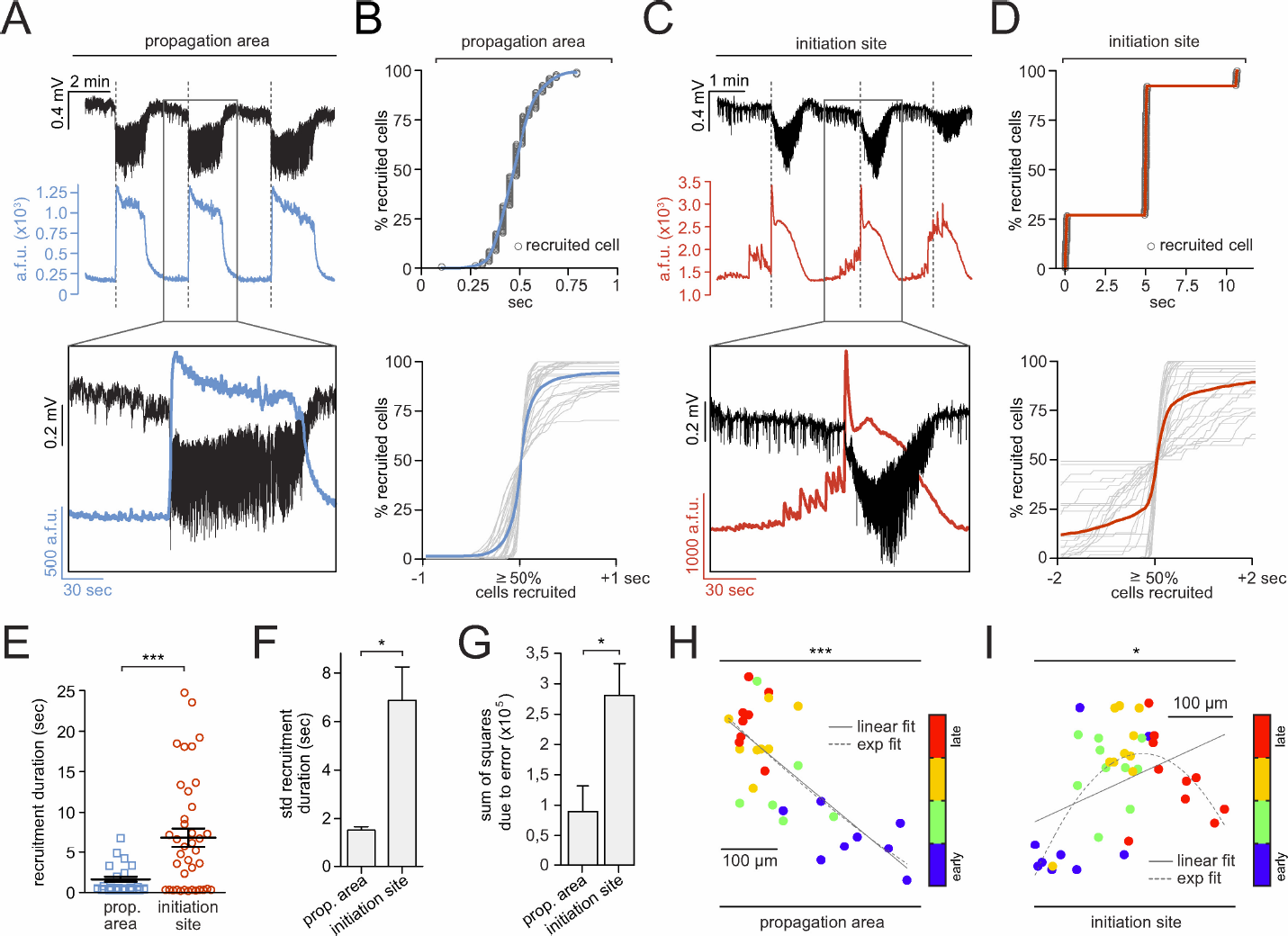
Differential intrafocal and extrafocal neuronal recruitment during epileptic. (**A**) Propagation area: LFP recording (black) and corresponding population average calcium transient (blue) of 3 consecutive seizures, detailed depiction of 2nd event (inset, suppl. movie 1). Note how electrographic seizure onsets (dotted lines) correspond to sudden rise of the population calcium signal. a.f.u. = arbitrary fluorescent units (**B**) Top: Representative example of smooth, s-shaped cell recruitment during an individual seizure onset. Bottom: Superposition of neural recruitment curves across all analyzed seizure onsets (gray) centered around the 50% recruitment frame; blue graph represents mean (n = 3 exp., total # of seizures = 31 [10,3 ± 0,88 s.e.m.], total # of cells analyzed = 626 [209 ± 44 s.e.m.], cell number in % for comparability across exp.). (**C**) Initiation site: LFP recording (black) and corresponding average population calcium transient (red) of 3 consecutive seizures, detailed depiction of 2nd event (inset, suppl. movie 2). Note temporal mismatch of electrographic seizure onsets and intrafocal population calcium events. Large pre-ictal population bursts are visible prior to the electrographic seizure onset (dotted line). (**D**) Top: Representative example of step-wise cell recruitment during intra- focal ictal progression. Bottom: Superposition of all analyzed intra-focal microseizures (gray) centered around the 50 % cell recruitment frame; red graph represents mean (3 exp., total # of seizures = 39 [13 ± 1,55 s.e.m.]). (**E**) Comparison of extra- and intra-focal cell recruitment durations (n = 31 and 39 durations; mean 1,83 ± 0,3 sec and 6,82 ± 1,1 sec, Mann Whitney test p<0,001). Determination of recruitment durations by calculating the time period from the first to the last recruited registered cell, excluding the 5% most deviant cells. (**F**) Comparison of extrafocal vs. intrafocal recruitment duration variability (mean std ± s.e.m., Mann Whitney test p<0,05). (**G**) Comparison of extrafocal vs. intrafocal seizure trajectories. Displayed sum of squares due to error based on a goodness of fit to a linear spatiotemporal ictal expansion. (**H**) Spatial analysis of propagation area: Spatiotemporal quartile clustering (quartiles calculated as mean coordinate of 1-25% earliest cells, 25-50%, 50-75% and 75-100%, see materials and methods) across 10 consecutive seizures (bivariate ANOVA p<0,001, all extrafocal exp. p<0,05). (**I**) Spatial analysis of initiation site: Spatiotemporal quartile clustering (each quartile coordinate = spatial mean of 25% recruited cells) clustering across 10 consecutive seizures (bivariate ANOVA p=0,0145, all intrafocal exp. p<0,05). Note how ictal progression in the propagation area and seizure initiation site visibly follow differential spatiotemporal trajectories (linear and exponential trajectory fit indicated by continuous and dotted lines).

The seizure initiation site, however, displayed strikingly different activity patterns (Figure 2 C). Unlike the surrounding cortex, we observed locally restricted, pronounced pre-ictal bursts of calcium activity, sometimes long before electrographic seizure onset (Fig. 2 C, suppl. movie 2). These neuronal coactivations (or ensembles) are consistent with previous observations of “microseizures” that can precede macro-electrographic seizures by seconds to minutes (Goldensohn, 1975; Schevon et al., 2008; Stead et al., 2010), or even weeks to months during epileptogenesis (Bragin et al., 2000). Consequently, the terms “ictal” and “pre-ictal” become somewhat blurred so we refer to the seizure onset as the electrographic (LFP) seizure onset. Moreover, neurons within the epileptic focus were recruited in a stepwise pattern (Fig. 2 D top). Indeed, when we superimposed all imaged intrafocal neuronal activity around the 50% recruitment frame, a heterogeneous pattern emerged from highly variable individual recruitment graphs (Fig. 2 D bottom, n = 3 experiments, GCaMP6s, total # of seizures = 39, average # of seizures 13 ± 1,55, total # of analyzed cells = 272, average # of analyzed cells = 92 ± 23 cells). Neuronal recruitment was temporally stretched, up to dozens of seconds ahead of the electrographic seizure onset (Figure 2 E), much beyond the already considerable variability of neuronal activation across seizures within the surround cortex (Wenzel et al., 2017) (Figure 2 F).

We compared the spatiotemporal maps of successive seizures between the propagation area and the seizure initiation site and found that both territories followed differential trajectories during ictal progression. Within the propagation area, ictal expansion followed a linear path (Figure, 2 G and H, suppl. movie 1), whereas in the initiation site, it was multi-directional (Fig. 2 G and I, suppl. movie 2). However, despite the differential trajectories, a conserved spatial pattern of relative cell recruitment was evident in both compartments across seizures (Figure 2 H and I, suppl. Fig. 1 A and B) (Neumann et al., 2017; Trevelyan et al., 2006; Truccolo et al., 2014; Wenzel et al., 2017). To quantify and compare these observed spatiotemporal patterns, we used a 2-dimensional ANOVA (Wenzel et al., 2017), categorizing cells into temporal quartiles and comparing i) the variance of the distance of each cell to the quartile spatial mean to ii) the variance of the distance to the spatial mean of all cells (Suppl. Fig. 1 C). The analysis yielded significant differences between distributions, with bivariate F-values for all experiments in the propagation area (F = 12,67, 11,64, 41,22; all p < 0,001) and in the epileptic focus (F = 22,15, 4,02, 3,43; p = 1,4×10-9, 0,0145, 0,024) (Fig. 2 H and I).

Taken together, we find a saltatory, multi-directional micro-epileptic seizure expansion within the epileptic focus followed by a smooth, more linear invasion of surround cortex. Within the epileptic focus, spatially restricted synchronizations occur up to dozens of seconds prior to electrographic seizure onset, with highly variable, temporally stretched neural recruitment time courses. In stark contrast, in the surround cortex, pre-ictal activity is low and ictal recruitment occurs only during full electrographic seizures. Despite differential recruitment time courses and spatiotemporal trajectories, the relative spatiotemporal activity patterns are consistent across seizures in both intrafocal and extrafocal compartments.

### Pre-ictal subnetwork compartmentalization and critical state transitions

The differential intrafocal and surround sub-network activity before electrographic seizure onset prompted us to further investigate the mechanisms underlying these pre-ictal population dynamics. To this end, we analyzed neuronal correlations in the calcium imaging data by applying principal components analysis (PCA; materials and methods) to 40-second pre-seizure periods. PCA weights were calculated within an experiment across all seizures and applied in time to simplify the “state-space” of multidimensional population activity occurring before and during seizure onset (Fig. 3 A and B). We found a multitude of substantial pre-ictal network state trajectories (Fig. 3 A, gray scale) within the epileptic focus prior to seizure onset (green). Indeed, compared to the pre-ictal network activity in surrounding cortex (Fig. 3 B, gray scale, onset in green), which was sparse (Schevon et al., 2012) and correspondingly did not show significant excursions in state space before seizure onset, intra-focal pre-ictal dynamics showed large epileptiform trajectories in PCA state space. During the 40 seconds before seizure onset, intra- focal average population activity was persistently higher than during baseline and escalated noticeably towards seizure onset (Fig. 3 C, red trace). On the contrary, pre-ictal population activity in the surrounding territories was in fact steadily lower than during baseline conditions (Fig. 3 C, blue trace).

**Figure 3:**
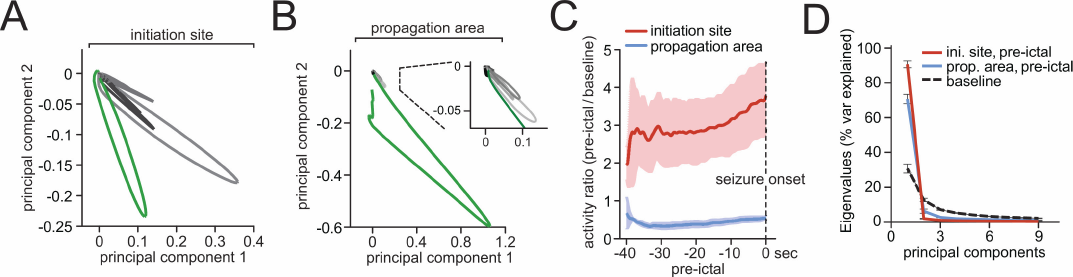
**Differential dynamical trajectories in intrafocal and extrafocal areas** (**A**) Representative network state spaces (40 second pre-ictal state changes in gray-scale [black = earliest change], electrographic seizure onset in green) within epileptic focus and (**B**) propagation area, with a magnified inset. (**C**) Cumulative frame-population-averaged positive change of fluorescence, normalized to baseline (set to 1), as indicator of general relative network activity changes over time (GCaMP6s, 3 extrafocal / 3 intrafocal experiments, 29 / 39 pre-ictal periods). Pre-ictal intra-focal versus penumbral network activity was compared by a running t-test (p<0.001 from −34 to 0 sec). (**D**) Scree plot of principal components of baseline network activity versus pre-ictal intrafocal and extrafocal activity. Note reduced number of principal components in both pre-ictal conditions. Displayed components accounted for more than 90% of the variance, respectively.

Interestingly, pre-ictal PCA trajectories were not simply lower magnitude versions of the ictal ones but occupied different portions of the multicellular state-space, suggesting that pre-ictal activity within the epileptic focus may involve specific subgroups of neurons rather than lower magnitude pulsing of the whole ictal population. A similar phenomenon was recognizable in the propagation area as well (Fig. 3 B, magnified portion, suppl. Fig. 2), where, during baseline conditions, a much greater diversity of network states emerged, stretching out in multiple dimensions (Fig. 3 B, magnified portion, suppl. Fig. 2 A). Thus, the transition from a pre-ictal to an ictal state involved a reduced dimensionality in population activity (Fig. 3 E). Such a transition suggests a decline of the network into lower dimensional attractors (or semi-stable, theoretically recurrent population states), which has been proposed by computational simulations and EEG analysis (Lopes da Silva et al., 2003) to occur during ictal transition, but never directly documented at cellular resolution.

Together, we show that during epileptic expansion, intra- and extrafocal subnetworks of neurons activate differently. While intrafocal population activity escalates towards electrographic seizure onset, surround cortical subnetwork activity drops below baseline level. In both compartments, subnetwork activity declines into lower dimensional attractors during transition to macroseizures.

### Local PV interneuron populations enhance ictal network compartmentalization

Recent *in vitro* and *in vivo* studies involving functional interference with interneuronal subtypes (especially fast-spiking PVs) have led to controversial results regarding the role of fast-spiking interneurons in seizure promotion or restraint (Avoli and de Curtis, 2011; Avoli et al., 1993; Cammarota et al., 2013; Gnatkovsky et al., 2008; Krook-Magnuson et al., 2013; Ledri et al., 2014; Shiri et al., 2015). With regard to interneuronal recruitment dynamics during epileptic activity, a recent study showed reliable recruitment of putative fast-spiking interneurons to epileptic activity (Neumann et al., 2017) echoing the finding of reliable ictal recruitment reliability at the general population level (Truccolo et al., 2011; Wenzel et al., 2017). In line with Muldoon and colleagues (Muldoon et al., 2015), Neumann et al. also found that epileptic activity predominantly entrained PVs, not pyramidal cells. However, it remains unclear, whether these studies recorded inside the seizure initiation site, or surround cortex.

To understand the role of local GABAergic interneuron populations during seizure formation and spread, we studied calcium dynamics of PV neurons in both intrafocal and extrafocal compartments by using transgenic mice that express the red fluorescent protein td-Tomato in parvalbumin containing interneurons (Madisen et al., 2010). First, we simultaneously imaged PV and non-PV calcium dynamics during seizure spread into the propagation area (Extrafocal, Figure 4 A, suppl. movie 3). Interestingly, PVs were consistently among the neurons displaying the strongest calcium activity during the pre-ictal period (Fig. 4 B). Population level analyses further supported this finding. While absolute activity (e.g. firing rates) could not be directly compared between PV cells and non-PV cells given known differences in cell calcium dynamics, bursting patterns, and baseline rates (Hofer et al., 2011), the relative ratio of PV versus non-PV population calcium transients was inverted in the pre-ictal period compared to baseline conditions (Fig. 4 C), suggesting that PVs comprise a functionally distinct sub-population in the propagation area during seizure formation (Liou, 2018; Neumann et al., 2017). During the initial phase of electrographic seizures, we encountered surprisingly eclectic PV population dynamics with several striking features. In accordance with previous reports, we identified PVs that reliably showed strong calcium transients just ahead of the arriving ictal wavefront (Fig. 4 D cell 1 and 3) (Cammarota et al., 2013; Kawaguchi, 2001; Schevon et al., 2012; Timofeev et al., 2002; Trevelyan et al., 2006; Ziburkus et al., 2006). At the same time however, we found at times immediately neighboring PVs displaying delayed recruitment (Cell 2 and 5 in Fig. 4 D and E). In line with the electrophysiological data by Neumann and colleagues (Neumann et al., 2017), these sequential PV calcium dynamics were conserved across seizures (Fig. 4 E). Surprisingly, we also identified transiently non-participant PVs (Cell 4 in Fig. 4 D upper panel, Fig. 4 E, suppl. movie 3.1). Clear recruitment to later events (Cell 4 in Fig. 4 D, lower panel, Figure 4 E, suppl. movie 3.2) excluded the possibility of a lack of GCaMP expression.

**Figure 4:**
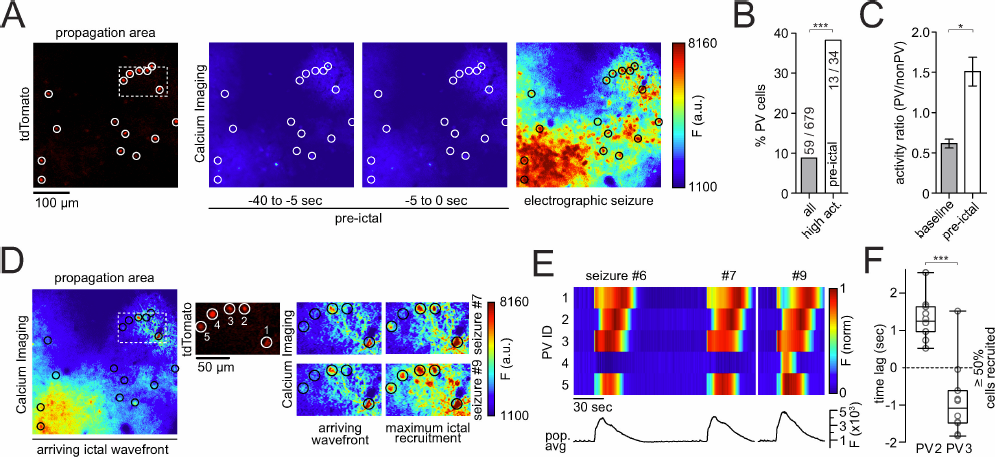
**Interneuronal recruitment in the propagation area** (**A**) Left to right: tdTomato-positive parvalbumin containing interneurons (PVs, encircled), imaged at 990 nm; calcium imaging (at 940 nm): fluorescence average images during pre-ictal period and sustained ictal activity. (**B**) PV percentage among highest active neurons during pre-ictal period (n = 4 experiments, total # of pre-ictal periods = 30 [7,5 ± 1,85 s.e.m.], all imaged non-PVs = 620, PVs = 59, top 5% activity cells = 34) as compared to general subtype distribution (Chi Square test P(χ2 >39,381) <0,001). (**C**) Activity ratio of PV versus non-PV sub- populations during baseline and the 40-second period before seizure onset. Activity ratios were obtained by dividing frame-sub-population-averaged positive change of calcium fluorescence. During baseline (left), PVs continually display lower values than non-PVs. During the pre-ictal period (right), this relationship is inverted (n = 4 experiments, GCaMP6s, 30 pre-ictal periods, Mann Whitney test p=0,0286). (**D**) Same experiment as in A, during break-in of ictal wave front (left); middle and right side: insert shown in A and D (dotted box). Displayed are 5 immediately neighboring PVs and their calcium transients during ictal break-in and sustained ictal activity across two successive seizures (upper and lower panel, see also suppl. movie 3.1 and 3.2). Note the diverse PV recruitment to ictal activity with PV #1 and #3 bursting ahead of the arriving ictal wavefront, PV #2 and #5 showing delayed recruitment (intra-ictal section), and one non- participant PV (#4) during the first seizure. The latter cell is clearly recruited to a successive ictal event. (**E**) Bottom: Population average calcium fluorescence signal of all 124 registered cells from exp. shown in A and D across three seizures (#6, 7,and 9). Top: Max-normalized calcium fluorescence for the 5 PVs highlighted in D, shown across three seizures. Note the differential recruitment on the order of seconds of even immediately neighboring PVs. PV #4 does only show a clear calcium signal in the last seizure displayed (**F**) Differential relative recruitment (Time lag with respect to the 50% recruitment frame) of immediately neighboring PV #2 and #3 (inter-cell distance <50 µm, both cells are located far from the arriving wavefront within the imaged area) across all seizures of an experiment (10 seizures, Mann Whitney test p<0,001). Boxes represent 25%ile to 75%ile of cellular recruitment, bands inside boxes display median cell recruitment time points, circles represent cell recruitment time lag for individual seizures.

Finally, we went on to image sub-population dynamics within the epileptic focus (Fig. 5 A, suppl. movie 4). Consistent with our results in the propagation area, PVs showed less population average calcium activity than non-PVs at baseline (Fig. 5 B left). Yet, by contrast, after 4-AP injection, this relationship was not inverted in the 40 seconds before seizure onset (Fig. 5 B right). We also did not encounter an increased percentage of intrafocal PVs among the neurons displaying highest calcium activity during the pre-ictal period. This does not indicate that intra- focal PVs were less active but, rather, that they were confronted with enhanced local firing of non-PV cells (Fig. 5 C). Like in the propagation area, we found diverse local recruitment of PVs to ictal activity (Fig. 5 D, arrowheads). During intra-focal micro-epileptic progression, we observed stepwise failure of the inhibitory surround (Cammarota et al., 2013; Trevelyan et al., 2006) (Figure 5 A and D, Fig. 5 D, blue bars, suppl. movie 4.1 and 4.2).

**Figure 5:**
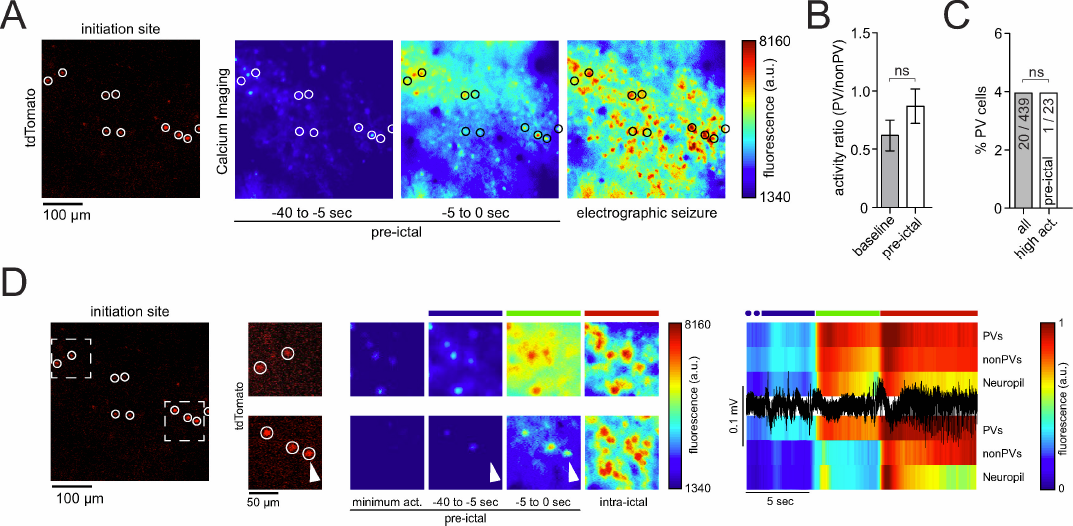
**Interneuronal recruitment inside the epileptic focus** (**A**) Left: tdTomato-positive parvalbumin containing interneurons (PVs, red, encircled) imaged at 990 nm; calcium imaging (at 940 nm): fluorescence average images of pre-ictal and intra-ictal neural activity (see also suppl. movie 4). (**B**) Activity ratio of PV versus non-PV sub-populations during baseline and the 40-second period before seizure onset. Activity ratios were obtained by dividing frame-sub-population-averaged positive change of ΔF/F (n = 3 experiments, all imaged non-PVs = 439, all PVs = 20, 23 pre-ictal periods). During baseline (left), PVs continually display lower values than non-PVs. This relationship is preserved during the pre-ictal period (right, Mann Whitney test p=0,4). (**C**) PV percentage among highest active neurons during pre-ictal period (n = 3 experiments, 23 pre-ictal periods, top 5% activity cells = 23) as compared to general subtype distribution (Chi-Square test P(χ2 > 0.002) =1,0151). (**D**) Same experiment as in A with two magnified inserts (see also suppl. movie 4.1 and 4.2) within distinct sub-territories. Calcium imaging during saltatory micro-epileptic progression (middle); note again the diverse recruitment of PVs to ictal activity (arrowhead). Colored bars above the calcium average images correspond to the colored bars above the LFP trace (black) and population average calcium traces (PVs, non-PVs and Neuropil) on the right.

We conclude that PVs have heterogeneous spatiotemporal patterns of activity within epileptic networks. Before electrographic seizure onset, a hypoactive surround cortex, and even small hypoactive regions (<100µm^2^) within the seizure initiation site, are observed together with increased local PV population activity. On a population level, PVs increase their firing ahead of the ictal wave front during ictal spread, in line with previous studies (Cammarota et al., 2013; Gnatkovsky et al., 2008; Prince and Wilder, 1967; Schwartz and Bonhoeffer, 2001; Timofeev et al., 2002). However, at the single cell level, we find side-by-side early and late PV recruitment in a spatially heterogeneous fashion, and even non-participant PVs. The temporal patterns of sequential PV activation repeat across ictal events (Neumann et al., 2017).

## Discussion

To investigate how epileptic seizures start within epileptic foci, we used an *in vivo* mouse model of acute pharmacological seizures induced by local cortical 4-AP injection (Liou, 2018; Ma et al., 2013; Wenzel et al., 2017; Zhao et al., 2011), a widely established model of partial onset neocortical seizures that enables the establishment of a territorially restricted epileptic focus whose location is precisely defined. We used fast calcium imaging to study cortical circuit activity within the epileptic focus and in primarily unperturbed surround cortex during secondary seizure spread.

We find intra-focal synchronization of neuronal populations that occurs *before* electrographic seizure detection (Figure 2). These coactive neuronal ensembles likely correspond to the microseizures described in human recordings (Goldensohn, 1975; Schevon et al., 2008; Stead et al., 2010; Worrell et al., 2008). Studies using multi-electrode arrays (MEA) have suggested that sustained micro-epileptic activity may represent the earliest step during seizure emergence (Bragin et al., 2000; Goldensohn, 1975; Schevon et al., 2008; Stead et al., 2010; Worrell et al., 2008). However, it has been difficult to definitely prove this hypothesis using MEAs, due to their sparse sampling of cortical circuits. Indeed, common MEAs used in humans cover a cortical area of 4×4 mm and contain 96 electrodes spaced 400 µm apart. Since the seizure initiation site can be as small as 0.04 mm^3^ and the exact locus of the seizure onset zone is usually poorly mapped in human patients or animal models of chronic epilepsy, full temporal seizure evolution is likely missed (Schevon et al., 2008). Our experiments, using an imaging method where we can monitor the activity of every neuron in the field of view, are consistent with the hypothesis that microepileptic activity is a prerequisite of seizures, as in our data ictal events were always reliably preceded by pre-ictal micro-epileptic build-up within the epileptic focus.

Our imaging approach further enabled us to map seizure progression from its earliest time point in the intact brain. We observed differential modes of spatiotemporal intrafocal progression during microseizures in comparison to ictal invasion of neighboring cortex during electrographic seizures. Inside the epileptic focus, we find a saltatory (i.e. “stepwise”, “modular” or “discontinuous”) expansion of ictal activity. These saltatory patterns are different to those observed in brain slices when applying disinhibitory drugs such as picrotoxin or bicuculline (Adams et al., 2015; Wadman and Gutnick, 1993). While there, “saltatory” propagation was observed on a millisecond range, our results show progression on a time scale of seconds, consistent with previous *in vitro* work where inhibitory circuits were kept intact (Trevelyan et al., 2006). This saltatory intrafocal activity during microseizures was followed by a more continuous seizure spread into nearby cortex during electrographic seizures, which suggests that seizure evolution consists of different consecutive steps of progression across anatomical scales (Figure 6, schematic overview). Smooth spread of ictal activity into nearby cortex surrounding an epileptic focus has been described at the macroscale (Ma et al., 2013; Schwartz and Bonhoeffer, 2001). Studies in human epilepsy patients using MEAs that were situated close, but not precisely inside the epileptic focus reported a continuous ictal progression as well (Schevon et al., 2012). Our results unify the different reported modes of ictal progression into one larger framework, and underscore that the precise location of recording is critical to any one experiment investigating progression of epileptic activity *in vivo*, not the least due to the pronounced spatial undersampling of current high resolution recording techniques such as MEAs or two-photon calcium imaging. Observed modes of progression likely depend on the experimental approach, investigated anatomical scale, recording location, recording modality and spatiotemporal resolution.

**Figure 6:**
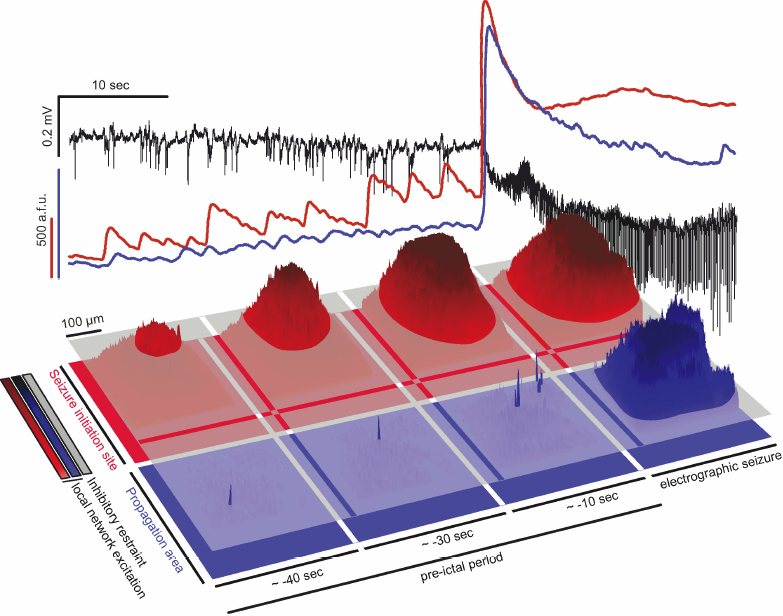
**Two-step seizure progression model** Top panel: Depicted are 40 seconds of the pre-ictal period prior to electrographic seizure onset, and initial 20 seconds of a full ictal event. LFP (black), and corresponding intrafocal (red) and extrafocal (blue) average population calcium transient. Note how during the pre-ictal period, increasingly escalating population calcium events are only detected inside the epileptic focus, not in extrafocal territories. Lower panel: Corresponding to the population average signals shown above, 3-D surface calcium activity plots of imaged fields of view inside the epileptic focus (red) and within extrafocal territory (blue). The gray layer schematically represents local interneuron population activity. During microseizures in the pre-ictal period, inhibition fails at the level of local ensembles only inside the epileptic focus. Small patches of excitatory ensemble activity break through the layer of local inhibition. Micro-epileptic expansion over the 40-second pre-ictal period occurs in a saltatory, non-linear fashion, with increasingly large areas where inhibition has failed to restrain epileptic activity. This expansion happens in the absence of an ictal LFP signature. Once a local threshold is reached (whose nature remains unclear), microseizures start spreading into neighboring territories outside the epileptic focus during electrographically detectable seizures, in a continuous fashion. Prior to this macro-epileptic expansion during electrographic seizures, little population activity can be detected in surround cortex, as opposed to intrafocal territories. This is due to increased feed-forward inhibition in extrafocal areas that is driven by pathological microepileptic activity inside the epileptic focus. Thus, seizure progression consists of consecutive steps, and may display differential spatiotemporal local network and sub-population dynamics highly depending on the localization of recording.

Even though abnormal interaction of excitatory and inhibitory neurons within epileptic networks has been shown to play a pivotal role in the progression of epileptiform activity (reviewed in (Trevelyan and Schevon, 2013)), little is known about their fine scale *in vivo* sub-population dynamics within ictal foci. Our data indicates that the stepwise intrafocal progression could be related to a failure of local PV populations. During microseizures, at times dozens of seconds before the electrographic seizure onset in our experimental setup, the epileptic focus could be divided into small micropatches that became successively, and abruptly, invaded over the course of numerous pre-ictal barrages. Intriguingly, the yet to be invaded patches had highly active PV cells, with little activity of immediately surrounding non-PVs until ictal break-in. This fits well with previous *in vitro* studies that suggested that interneuronal depolarization block (Sessolo et al., 2015; Ziburkus et al., 2006), or a depletion of releasable GABA, cause inhibitory failure during ictal events. Notably, it never occurred in our experiments that all PVs in the field of view (∼400×400 µm) showed simultaneous pre-ictal calcium transients. Yet, based on optogenetic activation paradigms that included the simultaneous activation of PVs in large areas or even entire brain slices (Shiri et al., 2015), recent *in vitro* studies suggested that PVs actively entrain ictal network synchronization. Our results indicate that this elicitation of ictal discharges might not reflect the *in vivo* situation. However, sustained interneuronal activity may support network synchronization by rebound excitation upon inhibitory failure (Grenier et al., 1998; Sessolo et al., 2015).

How could the propagation area experience a different kind of ictal invasion, as compared to the epileptic focus? The reason for consecutive steps of seizure progression could lie in differential pre-ictal sub-network dynamics within intra- and extrafocal cortical compartments. We show that, while the seizure initiation site is experiencing a buildup of activity with locally restrained population bursts, the activity in the surround cortex drops below baseline levels, with simultaneously enhanced calcium dynamics of local PV populations. This is in line with previous literature that suggests that feedforward inhibition, driven by the ictal core via short- and long-range axonal projections, could place areas ahead of the ictal wavefront into an “ictal penumbra state” (Schevon et al., 2012; Trevelyan et al., 2007). This penumbra state seems to be present even at large distances to the focus during ongoing local seizure activity (Liou, 2018). Based on this, our result of enhanced PV population activity in cortex nearby the epileptic focus could account for the ictal progression we observe. The level of inhibition needed to restrain pathologically high levels of excitatory synaptic inputs into surround cortex prior to seizure generalization is presumably lower than in the epileptic focus itself. Thus, driven by the ongoing ictal buildup within the epileptic focus, PVs in surround cortex could suppress local network activity (which is what we observe in the propagation area), but thereby also slowly start an eventually detrimental process that has taken place already within the epileptic focus before seizure spread. This process could be driven by compartmentalized (e.g. epileptic focus versus surround cortex) extracellular and intracellular ionic disturbances such as intra-neuronal chloride accumulation (Alfonsa et al., 2015; Huberfeld et al., 2007; Pallud et al., 2014). A slow shift in the GABAergic reversal potential in surround cortex during ongoing activity build-up in the seizure initiation site could lead to a more rapid nearby surround cortex recruitment during seizure generalization, resulting in the observed, smoother ictal expansion. Of note, despite pre-ictal network compartmentalization (Prince and Wilder, 1967; Schevon et al., 2012; Schwartz and Bonhoeffer, 2001), both compartments, the seizure initiation site and the propagation area, share a dimensionality reduction of possible network states starting from early on during ictogenesis. In line with Liou and colleagues (Liou, 2018), our sub-population experiments indicate that this decline into lower dimensional attractors could be related to enhanced levels of widespread inhibition, which in turn might carry implications for the early detectionability of pending macroseizures in humans. The identification of such attractor changes would not require high spatial resolution recordings and could open up an extended pre-ictal time window for therapeutic intervention.

Finally, we find that the recruitment dynamics of local PV populations to seizure break-ins are spatially diverse, beyond the model of previous studies showing that interneurons fire ahead of the propagating ictal wave front (Cammarota et al., 2013; Kawaguchi, 2001; Schevon et al., 2012; Timofeev et al., 2002; Trevelyan et al., 2006; Ziburkus et al., 2006). We find a range of immediate to delayed full recruitment of local PV populations to cortical seizures, and that the temporal pattern of PV recruitment is conserved across ictal events. This is in line with Neumann and colleagues who performed tetrode recordings in chronically epileptic rats, and showed that putative fast-spiking interneurons were recruited to seizure activity in sequences preserved across ictal events (Neumann et al., 2017). This result is also complementary to our and other research groups’ recent analyses of local ictal network recruitment patterns (Rossi et al., 2017; Truccolo et al., 2014; Wenzel et al., 2017). Intriguingly, we also found non-participant PVs during full ictal events, which would be impossible to uncover using micro-electrode arrays, as silent neurons remain invisible to extracellular electrode recording techniques. Since PVs can synapse onto each other and are also targets of other interneuronal classes (Bezaire and Soltesz, 2013; Pfeffer et al., 2013; Pi et al., 2013; Sik et al., 1995), non-participant PVs, and potentially other inhibitory subtypes as well, may be actively inhibited during ictal activity (Paz and Huguenard, 2015). This finding might also help understand why in vivo optogenetic activation of PVs during seizures still suspended epileptic activity (Krook-Magnuson et al., 2013), in a circumstance where one would have assumed PVs to be strongly active anyway. In those experiments, light activation might have recruited predominantly “non-exhausted” PVs. What was surprising to us was that while temporal sequences of PV interneuron recruitment were reliable across seizures, their spatial recruitment on a single cell level was much more heterogeneous than previously thought. At times, immediately neighboring PVs (inter-cell distance < 20 µm) showed completely different temporal activation. This uncovers that below population level analysis (which relates to the average behavior of a group of cells), the PV sub- family does not activate homogeneously ahead of the traveling ictal wavefront. Instead, local ictal PV activity patterns are spatially diverse, potentially due to differential PV subtype connectivity patterns within this sub-family.

In summary, we provide novel insights into the microcircuit dynamics during across seizure evolution from its earliest time point in vivo. Our experiments help unify previously contrasting results on modes of ictal progression and show that ictal PV population dynamics in both surround and intrafocal regions are more diverse than previously thought. Future work on inhibitory, and dis-inhibitory activity at the microcircuit level within precisely defined locations across the epileptic network will aid in the development of more targeted and efficient future therapeutic routes.

**Acknowledgments:** We thank Dr. Yeonsook Shin, Alexa Semonche, Reka Recinos, Mari Bando and Azadeh Hamzei for viral injections. Additionally, we are grateful to other members of the Yuste Lab for useful comments. This work was supported by the Deutsche Forschungsgemeinschaft (DFG, grant WE 5517/1-1), NEI (DP1EY024503, R01EY011787), NIMH (R01MH101218, R01 MH100561) and DARPA SIMPLEX N66001-15-C-4032. This material is based upon work supported by, or in part by, the U. S. Army Research Laboratory and the U. S. Army Research Office under contract number W911NF-12-1-0594 (MURI). The authors have no competing financial interests to declare. M.W. and R.Y. conceived the project. M.W. performed all experiments and wrote the paper. M.W. and J.P.H. analyzed the data. All authors planned experiments, discussed results and edited the paper. R.Y. assembled and directed the team and secured funding and resources.

## Methods

All experiments were performed with care and in compliance with the Columbia University institutional animal care guidelines. Experiments were carried out on either C57BL/6 wild type mice or PV-Cre::LSL-tdTomato mice at postnatal age of 1-3 months. Animals were never used for previous or subsequent experiments. Food and water was provided ad libitum. Mice were housed at a 12 hour light/dark cycle.

**Virus injections and surgical procedures.** Prior to actual experiments, animals were injected with AAV1-Syn-GCaMP6s (purchased from the University of Pennsylvania Vector Core). Mice were anesthetized with isoflurane (initial dose 2-3% partial pressure in air, then reduction to 1- 1.5%). A small cranial aperture was established above left somatosensory cortex (coordinates from bregma: x 2,5mm, y −0,24mm, z −0,2mm) using a dental drill. A glass capillary pulled to a sharp micropipette was stereotactically lowered into cortical layer 2/3. A 800nl solution of 1:1 diluted AAV1-Syn-GCaMP6s (Chen et al., 2013) was slowly injected over 5 min at a depth of 200 µm from the pial surface using a microinjector (World Precision Instruments). 4-5 weeks after virus injection, on the day of the experiment, mice were anesthetized with isoflurane (initial dose 2-3% partial pressure in air, then reduction to 1.0%). A small flap of skin above the skull was removed and a titanium head plate with a central foramen (7×7mm) was attached to the skull with dental cement above the left hemisphere (Fig 1a). Then, a small craniotomy similar to previous descriptions (Miller et al., 2014) was carried out. Specifically, posterior to the virus injection site, the skull was circularly thinned using a dental drill until a small piece (circa 2mm in diameter) of skull could be removed effortlessly with fine forceps.

**Two-photon calcium imaging.** Activity of cortical neurons was recorded by imaging changes of fluorescence with a two-photon microscope (Bruker; Billerica, MA) and a Ti:Sapphire laser (Chameleon Ultra II; Coherent) at 940 nm through a 25x objective (water immersion, N.A. 1,05, Olympus). Resonant galvanometer scanning and image acquisition (frame rate 30 fps, 512 x 512 pixels, 100-170 µm beneath the pial surface) were controlled by Prairie View Imaging software. Multiple datasets were acquired consecutively over the course of an experiment (90,000-150,000 frames in total, with several momentary breaks interspersed for reasons of practicality). During the entire experiment, the head-restrained animals were kept under light isoflurane anesthesia (0.8-1% partial pressure in air) via a nose piece while body temperature was maintained with a warming pad (37.5°C).

**Ictal model and Electrophysiology.** The local 4-AP model of acute cortical seizures was used in this study, as it allowed the precise localization (on the order of micrometers) of the epileptic focus and surround cortex. In the context of the model used here, it is further noteworthy that seizure occurrence in chronic animal models of epilepsy is usually low (Arida et al., 1999; Ewell et al., 2015; Muldoon et al., 2015). It would require either multiple fortunate or prohibitively long imaging sessions of infrequently occurring full ictal events in order to capture a sufficient number of seizures, along with reasons concerned with potential tissue photo-damage, fluophore bleaching and hardware limitations. For local field potential (LFP) recordings, a sharp glass micropipette (2-5 MΩ) filled with saline and containing a silver chloride wire was carefully advanced into the cortex (30° angle) under visual control to a depth of around 100 µm beneath the pial surface. The pipette tip was positioned close by the imaged area (Fig. 1 A). A reference electrode was positioned over the contralateral prefrontal cortex. LFP signals were amplified using a Multiclamp 700B amplifier (Axon Instruments, Sunnyvale, CA), low-pass filtered (300Hz, Multiclamp 700B commander software, Axon Instruments), digitized at 1 kHz (Bruker) and recorded using Prairie View Voltage Recording Software alongside with calcium imaging. For induction of ictal events, another sharp glass micropipette containing 4-Aminopyridine (4-AP, 15mM, 0.5 µl) was slowly lowered (30° angle) into the cortex to a depth of 420-480 µm. The pipette tip was positioned at a distance of around 1.5-2 mm caudally to the imaged area. Correct positioning of the pipette tip was ensured by a diagonal dry-run within saline above the cortex preceding actual insertion. 4-AP was injected over the course of 10-15 min by use of a Micro4 Micro Syringe Pump Controller (World Precision Instruments). Electrographic seizure onset time points were determined mathematically by mean and standard deviation (std) of LFP recordings. The first time point exceeding > 5 std from the interictal LFP mean power was defined as the seizure onset and confirmed by visual inspection.

**Image Analysis.** Cell regions of interest (ROIs) were identified in a semi-automated fashion by using custom written software in MATLAB (Caltracer 3 beta, available at our laboratory website: http://www.columbia.edu/cu/biology/faculty/yuste/methods.html) followed by manual confirmation. Because of pronounced and synchronous fluorescence changes of surround neuropil during epileptic conditions, halo subtraction procedures could lead to distortions of calcium transients of individual cells. Therefore, we applied ROI shrinkage (ROI soma outline minus 1.5 pixels radially), which has been successfully used to minimize bleed-in of surround neuropil fluorescence (Hofer et al., 2011). Cells with low signal to noise ratio or no apparent calcium transients were excluded from further analysis. Individual cells that were lost over the course of the experiment due to at times discrete axial movement of the imaged focal plane during local 4-AP injection were also excluded from further analysis. Individual cell fluorescence was calculated as the average across all pixels within the ROI.

**Cell Recruitment Analyses.** In order to determine the recruitment time-point of individual cells to ictal activity, we used the first discrete derivative (slope) of the individual fluorescent traces assuming that the sharpest change in fluorescence correlates best with maximal recruitment to ictal activity (Trevelyan et al., 2006; Wenzel et al., 2017). Population recruitment durations were calculated as the time from the first to the last recruited registered cell, excluding the 5% most deviant cells to reduce outlier-induced duration distortions. Specific time lags of PV neural recruitment in comparison to local network recruitment to seizure activity were calculated as follows (Suppl. Fig. 3). First, we derived the median frame *Y_50_* wherein the cumulative number of recruited cells first equaled or exceeded 50% of all registered neurons. Then, relative to this *Y_50_*, each PV was assigned a temporal recruitment lag by subtracting *Y_50_* from its individual recruitment frame *X_N_* (*X_n_ – Y_50_*, a negative result would indicate an early recruitment of the respective PV ahead of the median population recruitment for a given seizure event).

**Spatiotemporal Clustering.** In order to identify spatio-temporo-progressive network activity motifs during progression of ictal activity, we ordered cell recruitment points in time (from early to late recruitment) during each individual ictal event of an experiment and divided each dataset into four groups (1-25%, 26-50%, 51-75%, 76-100%), as previously described (Wenzel et al., 2017). Then, the mean distance of cell coordinates (x,y within imaged area) of individual quartiles were calculated to either the mean coordinate of the respective quartile, or the mean coordinate of all recorded cells, for each seizure. Then, these distances were compared to each other. If cells’ recruitment time points are similar and cells cluster spatially, their distance to the mean coordinate of their respective quartile should be significantly shorter than the mean distance to the mean coordinate of all cells which are distributed over the entire field of view (Suppl. Fig. 2 C).

**State Space.** We sought to parsimoniously describe multicellular network dynamics in a lower rank subspace amenable to visual display and linear comparisons. We calculated a principle components analysis (PCA) on i) baseline (pre-injection) activity, ii) 40 second period prior to seizure onset, and iii) 3 seconds after seizure onset. We compared scree plots to quantify the proportion of variance accounted for by each component (Cattell, 1966), comparing root eigenvalues between conditions to assess an indication of dataset dimensionality, and, for plotting purposes, carried out a VARIMAX rotation to limit solutions to 3-6 components (based on scree plot).

**Statistics.** All data were analyzed using custom written code in MATLAB (MathWorks, Inc). Error bars on bar plots and shaded areas in graph plots indicate s.e.m. To determine statistical significance for analyses regarding cell recruitment, Mann Whitney tests were applied unless stated otherwise. For statistical analysis of spatio-temporal clustering, we used bivariate ANOVA analysis of mean distance differences (df = 3). High activity subtype comparison (Fig. 4 and 5) was determined by Chi-Square test (1 degree of freedom [df]).

### Additional resources

MOCO (ImageJ plugin for motion correction) and Caltracer (extraction of calcium signals) are available online on the Yuste laboratory web page: http://www.columbia.edu/cu/biology/faculty/yuste/methods.html

## Resource Table

| **Experimental Models: mouse lines** |  |  |
| --- | --- | --- |
| C57BL/6 wildtype | Jackson Laboratory | RRID:IMSR_JAX:00064 |
| PV-Cre | Jackson Laboratory | RRID:IMSR_JAX:017320 |
| LSL-tdTomato | Jackson Laboratory | RRID:IMSR_JAX:007908 |
| **Chemicals (Pharmacological compounds)** |  |  |
| 4-Aminopyridine | Sigma-Aldrich | Cat # 275875; CAS:504-24-5 |
| **Recombinant DNA** |  |  |
| AAV1-Syn-GCaMP6s-WPRE-SV40 | U Penn Vector Core | Cat # AV-1-PV2824 |
| **Software and Algorithms** |  |  |
| ImageJ | https://imagej.nih.gov/ij/ |  |
| Moco | http://www.columbia.edu/cu/biology/faculty/yuste/methods.html |  |
| MATLAB(R2014b) | MathWorks |  |
| Caltracer3beta | http://www.columbia.edu/cu/biology/faculty/yuste/methods.html |  |
| Adobe Illustrator CS6 | Adobe |  |

